# Tests of hybridisation in *Tetragonula* stingless bees using multiple genetic markers

**DOI:** 10.1101/2020.03.08.982546

**Authors:** James P. Hereward, Tobias J. Smith, Ros Gloag, Dean R. Brookes, Gimme H. Walter

## Abstract

Discrepancies in mitochondrial and nuclear genetic data are often interpreted as evidence of hybridisation. We re-examined reports of hybridisation in three cryptic stingless bee species in the genus *Tetragonula* in South East Queensland, Australia (*T. carbonaria, T. davenporti*, and *T. hockingsi*). Previous studies on this group using microsatellite markers proposed that occasional hybrids are found. In contrast, we find that allele frequencies at neutral regions of the nuclear genome, both microsatellites and random *snps*, reliably separated the three species, and thus do not support hybridisation. We found no inter-species variation in PCR amplicons of the nuclear gene *EF1alpha*, but low and moderate species-specific polymorphisms in the nuclear gene *Opsin* and the mitochondrial *16S* respectively, with no cases of mito-nuclear discordance at these genes. We confirm that nuclear divergence between these species is low, based on 10-26kb of non-coding sequence flanking *EF1alpha* and *Opsin* (0.7-1% pairwise difference between species). However, we find mitogenomes to be far more diverged than nuclear genomes (21.6-23.6% pairwise difference between species). Based on these comprehensive analyses of multiple marker types, we conclude that there is no ongoing gene flow in the *Tetragonula* species of South East Queensland, despite their high morphological similarity to one another and the low nuclear divergence among them. The mitogenomes and draft nuclear genomes provided for these species will be a resource for further molecular studies on this group, which are important pollinators in Australian natural and agroecosystems.

## Introduction

Allopatry is key to speciation, but species boundaries may be tested when populations that diverged in allopatry come back into secondary contact through changes in habitat or climate, or as a result of anthropogenic movement over geographic boundaries (Sánchez-Guillén *et al*., 2013, Gloag *et al*. 2017; Brookes *et al*. 2020). The process of hybridisation is closely tied to the designation of species in population genetic terms (Paterson 1978). When species are considered conceptually as recombining gene pools (Ayala *et al*. 1994), hybridisation is the process of gene flow across two such species gene pools. High levels of hybridisation in local areas of overlap indicate that the species in question would be better viewed as subspecies, which, by definition, have independent geographical distributions but hybridise freely where and when they do overlap (Walter 2003, Ford 1974). Such situations are often seen when closely related lineages of the same species are brought back into sympatry (Feder *et al*. 2003), but the pattern is also sometimes seen in more distantly related lineages (e.g. Shipman *et al*. 2019). The frequency of hybridisation, or gene flow, between gene pools is therefore key to understanding the limits of a given species, and to our understanding of speciation.

Accurate assessment of the frequency of hybridisation under natural conditions is therefore the first step in shedding light on the process of speciation. The more recently two populations have diverged, the more likely they have not yet become separate species. When two subspecies come back into secondary contact, gene flow is generally re-established (Feder *et al*. 2003). Hybrid detection between close relatives is especially sensitive to the molecular methodologies used (e.g. number and type of markers), and more likely to be susceptible to sampling error because parent species are similar in appearance with respect to one another.

An Australian species complex of stingless bees (Meliponini) in the genus *Tetragonula* illustrates these challenges well. The genus *Tetragonula* includes three cryptic sister species (*T. carbonaria. T. hockingsi* and *T. davenporti*), indigenous to eastern Australia, that are part of a monophyletic clade known as the “Carbonaria group” (Franck *et al*., 2004). These bees can be kept readily in wooden hives, are popular pets, and are effective pollinators of some tropical and subtropical crops (Dyer *et al*. 2016, Heard 1994). This has led to many thousands of colonies being kept by beekeepers in Eastern Australia. Today, the distributions of all three species overlap in parts of south-east Queensland (Fig. 1). Beekeeping and other human activities, such as land use change, have greatly increased the overlap of their distributions in recent decades, particularly through the southward spread of *T. hockingsi*. Unfortunately, the historical distributions of the species are poorly known due to their cryptic morphology.

**Figure 1.**
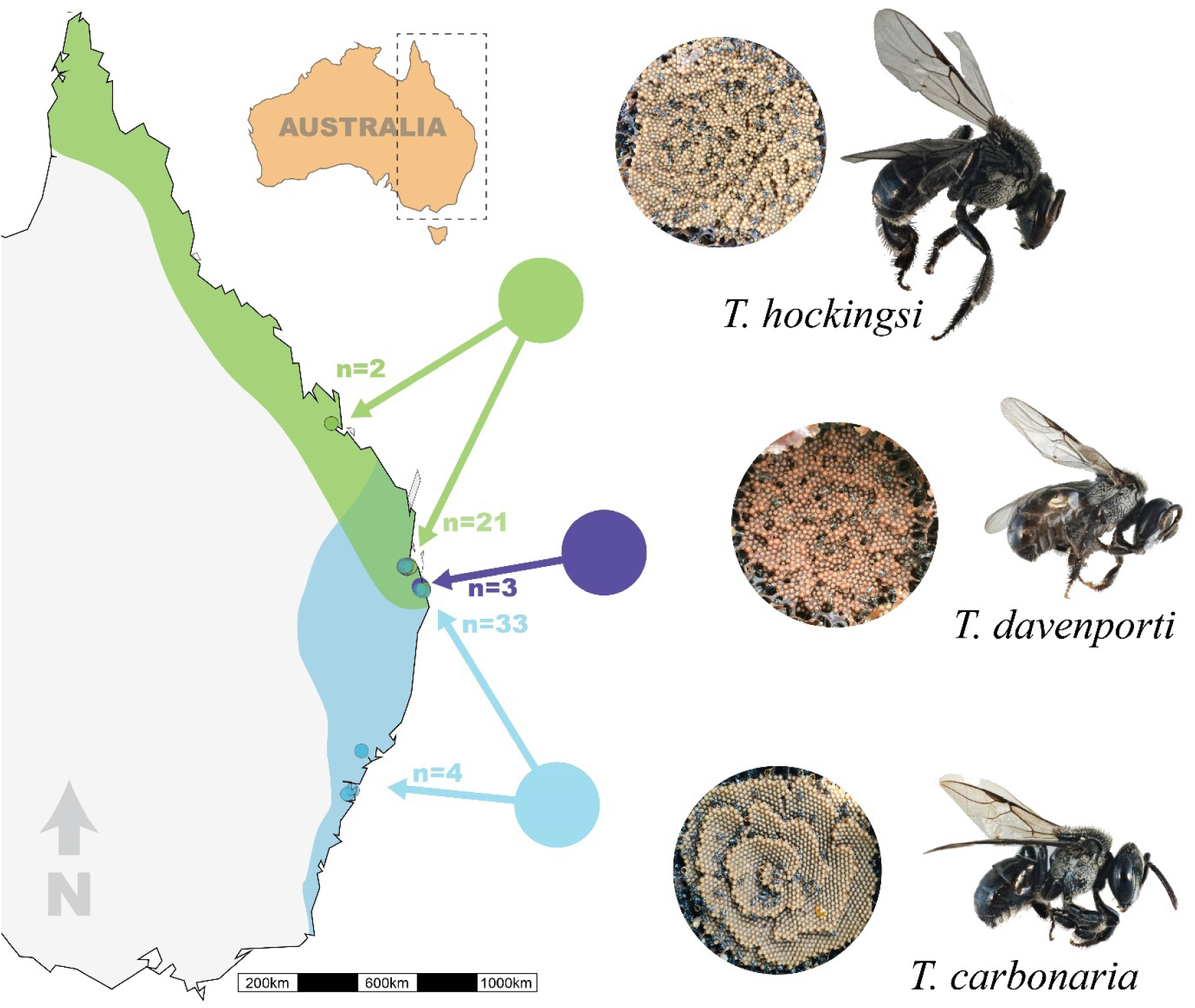
Map showing the known distributions of three species of *Tetragonula* in Australia, sampling sites are shown as small coloured circles, and numbers used in this study denoted by different colours for each species. The brood structure and the bees are shown on the right, and the distributions are based on Heard (2016).

Queens, males and workers (females) cannot be reliably identified based on morphology alone (Dollin *et al*. 1997). Rather, species are typically identified by the structure of their brood comb, with *T. carbonaria* producing a neat spiral of adjacent brood cells in a layered comb and *T. hockingsi* producing clusters of cells in a form known as semi-comb (Fig. 1). *Tetragonula davenporti* shares the semi-comb brood structure of *T. hockingsi*, and is known from only a handful of managed colonies in a sub-coastal pocket of southern Queensland (Franck *et al*. 2004, Brito *et al*. 2014), but is believed to persist in the wild in nearby forested areas (P. Davenport pers. Comm 2019; R. Gloag unpublished data).

Relatively little is known about the mating behaviour of these bees, because it occurs during the nuptial flight. Genetic data indicates that queen’s mate only once (Green & Oldroyd 2002), and observations of tethered virgins indicate that male genital detachment is likely the mechanism by which multiple mating is avoided (Smith 2020). Male mating congregations, which form outside colonies with virgin queens preparing for nuptial flights, can contain a mix of both *T. carbonaria* and *T. hockingsi* males (Cunningham *et al*. 2014). These two species are also known to engage in interspecific nest usurpations, in which an invading colony kills a resident queen and installs its own in the nest (Cunningham *et al*. 2014), a behaviour which may increase the chances that males are attracted to interspecific queens.

Previous efforts to establish if, and how regularly, gene flow occurs between these species, have returned equivocal results. First, microsatellite markers have been used to look for evidence of hybrids. Frank *et al*. (2004) identified some South-East Queensland colonies of *Tetragonula* as being of hybrid origin based on microsatellite data, where hybrids were defined as samples having a posterior probability of assignment of less than 90% to a cluster in an analysis using the genetic assignment program *STRUCTURE* (Pritchard *et al*. 2000). The same three Australian *Tetragonula* species were re-investigated by Brito *et al*. (2014), with microsatellite data and two mitochondrial genes. Three instances of hybridisation in South-East Queensland were reported within the zone in which *T. carbonaria* and *T. hockingsi* overlap. This was based on a threshold of 95% posterior probability of belonging to a cluster in a Bayesian clustering analysis of the microsatellite data. Brito *et al*. (2014) also included two hives reported to be *T. davenporti* in their study. One of the *T. davenporti* hives clustered with *T. carbonaria* in a Principal Coordinate Analysis (PCoA) of genetic distance based on the microsatellite data, and the other clustered slightly outside of the ‘*T. carbonaria*’ cluster. They suggested that *T. hockingsi* represents two genetic clusters, one each in northern and southern Queensland, based on their microsatellite and mitochondrial data. Their reported *T. hockingsi x T. carbonaria* hybrids were from South East Queensland and they suggested that this hybridisation was due to the anthropogenic movement of northern *T. hockingsi* into the southern region.

Second, both nuclear and mitochondrial markers have been used to look for evidence of introgression and thus past interbreeding across Australia’s East Coast *Tetragonula* species. These efforts were hampered by the amplification of nuclear copies of mitochondrial genes (numts) (e.g. for *cytochrome b* (*cytb*); Franck *et al*. 2004). Recently, several numts of Cytochrome Oxidase Subunit I (*COI*) were identified in *T. hockingsi* and *T. carbonaria* (Françoso *et al*. 2019), which cast into question even the designation of *T. davenporti* as a true species. Surprisingly, the genuine COI sequences of *T. hockingsi* and *T. carbonaria* are highly diverged (16.5 %), relative to available nuclear genes (Cunningham 2014, Francosco 2019). This pattern of high mitochondrial divergence but low nuclear divergence might be consistent with recent or ongoing gene flow and highlights the need for new analyses to understand the species status of these bees. This will in turn reveal the likely impact of continued movement of managed *Tetragonula* hives around Australia.

To resolve whether ongoing gene flow occurs between *T. carbonaria, T. hockingsi* and *T. davenporti* we sampled these species in sympatry in South East Queensland. We compared microsatellite data, using the same markers as previous studies, to a *genotyping by sequencing* approach that yielded 2,049 *snp* markers with less than 5% missing data. We also look for evidence of mitochondrial introgression, by considering two nuclear gene sequences and one mitochondrial gene sequence of each species. We then use Illumina sequencing to reconstruct mitochondrial genomes and confirm that our mitochondrial sequences are not pseudogenes. Finally, we assembled draft nuclear genomes for these three species as a resource for further genetic analysis of the species complex. This comprehensive approach leads to new insights about the extent of hybridisation in these bees, and their species status.

## Materials and Methods

### Sampling and DNA extraction

We sampled four workers per colony from hived colonies of each species between February 2017 and April 2019 in south-east Queensland (*T. carbonaria*: N=33, *T. hockingsi*, N=21, *T. davenporti*, N=3) and we also included two *Tetragonula hockingsi* hives from Rockhampton (Queensland), two hives of *Tetragonula carbonaria* from Newcastle (NSW), and two from Sydney (NSW) (Fig. 1). Most *T. carbonaria* and *T. hockingsi* colonies were sampled in the greater Brisbane region. We opened each hive and assigned it to species based on brood nest architecture (spiral/layered comb or semi-comb). The three *T. davenporti* colonies were sampled from the border area of Queensland and New South Wales and were the same colonies used for past genetic work on this species (P. Davenport, pers. comm. 2019). A further two *T. carbonaria* colonies and one *T. hockingsi* colony were sampled from the same meliponary as the three *T. davenporti* colonies. We extracted DNA using a silica spin-column method (Ridley *et al*. 2016) and normalized concentrations to 5ng/ul per sample following quantification (PicoGreen, Life Technologies, California, USA).

### Neutral nuclear markers

#### Microsatellites

We first assessed species status and evidence of hybridisation using seven microsatellite loci (Green *et al*. 2001), as used in previous molecular studies of this group (Franck *et al*. 2004, Brito *et al*. 2014). We used PCR reactions of 2 μl DNA template, 0.5 U MyTaq polymerase (Bioline, Australia), 0.1 μM of forward primer, 0.2 μM of reverse primer, 0.2 μM M13 labelled primer with different fluorescent dyes (6-FAM, VIC, PET or NED) and 1x buffer, with reaction conditions of an initial denaturation at 95 °C for 3 min, followed by 10 cycles of 10 s at 95 °C, annealing at 45 °C for 25 s, and 1 min extension at 72 °C, then 40 cycles of 10 s at 95 °C, annealing at 55 or 52°C for 25 s, and 1 min extension at 72 °C, and the final extension was at 72 °C for 2 min (see Cunningham *et al*. (2014) for locus-specific annealing temperatures). Loci were then pooled with one locus per dye, cleaned with 1U Exonuclease I and one unit of Antarctic Phosphatase, as above, and genotyped by Macrogen Inc. (Seoul, Republic of Korea). Microsatellite peaks were analysed using the microsatellite plugin in Geneious v. 2019.2.1 (http://www.geneious.com, Kearse *et al*. (2012)).

#### Single nucleotide polymorphisms

We generated genome-wide *snp* data using a genotyping-by-sequencing method. We adapted a protocol and adaptor regime based on the methods of Elshire *et al*. (2011), Poland *et al*. (2012) and Petersen *et al*. (2012), with barcodes based on Caporaso *et al*. (2012). Full details of the method are provided in supplementary data and can be found online at http://www.jameshereward.org/GBS.html. We pooled 288 individuals per sequencing lane, and sequenced the libraries with PE150 Illumina sequencing at Novogene (Beijing, China).

The sequence data were demultiplexed, assembled, and *snps* called using STACKS (Catchen *et al*. 2013). We then filtered the vcf using vcftools, with the data first being filtered to a minor allele count of three (one heterozygote and one homozygote), which allows the conservative removal of singleton *snps* that are likely to be errors, without discarding rare alleles (Linck & Battey 2019). We set a minimum depth of 5 and kept only biallelic *snps*. We filtered the data for missing data in three steps. First, any marker missing more than 50% data was discarded (i.e. 50% of individuals were not genotyped at that marker), to remove the markers most affected by missing data. Second, any individual that had missing data at more than 50% of the markers was discarded, to remove the individuals that had bad quality genotyping. Finally, any marker missing more than 5% data was discarded to produce a final dataset with relatively little missing data (∼3%).

For both our microsatellite and *snp* datasets, we performed principal component analyses (PCA) using the adegenet package in R (Jombart, 2008) and *STRUCTURE* analyses (Pritchard 2000). For the PCA we used the full *snp* dataset (2,049 *snps*), but for *STRUCTURE* only one *snp* per locus was retained (1,935 *snps*). *STRUCTURE* is an individual-based clustering algorithm that assigns individuals to each of *K* population clusters using Hardy-Weinberg equilibrium and linkage information in genetic markers. To assess the effect of replication and missing data on hybridisation inferences from the *STRUCTURE* output, we repeated the analyses on three versions of our microsatellite and *snp* datasets : (i) a dataset including one worker per colony, (ii) a dataset including all four workers per colony, and (iii) a dataset with 20% randomly deleted. To achieve 20% missing data in the the microsatellite dataset, data was removed evenly across markers until a 20% reduction was achieved. For the *snp* dataset, we changed the missing data threshold in the filtering step to 20%, and this increased the number of markers to 13,409 loci. Having more markers than the original dataset clearly makes for an uneven comparison, so we simply deleted every marker after the first 1,935 *snp* markers to make this dataset the same size as the original one with no missing data. For each *STRUCTURE* run, we used the admixture model and ran 500,000 iterations of burn-in followed by 1 million iterations. We assumed both two or three populations (K=2 and K=3) and performed multiple runs that were then permuted and plotted using Clumpak server (Kopelman *et al*. 2015).

### Gene sequencing

To confirm whether common marker genes showed species-specific polymorphisms, we sequenced two nuclear genes (*EF1alpha* and *Opsin*) and one mitochondrial gene (*16S*). These genes were selected as they have previously been used for a phylogenetic study of stingless bees (Rasmussen and Cameron, 2009). For *EF1alpha* we used primers F2-ForH (GGRCAYAGAGATTTCATCAAGAAC) and F2-RevH2 (TTGCAAAGCTTCRKGATGCATTT) (Hines 2006). For *Opsin* we used LWRhF (AATTGCTATTAYGARACNTGGGT) and LWRhR (ATATGGAGTCCANGCCATRAACCA) (Mardulyn & Cameron 1999). For *16S* we used primers 874-16S1R (TATAGATAGAAACCAAYCTG) (Cameron 1992) and 16SWb (CACCTGTTTATCAAAAACAT) (Dowton & Austin 1992). We performed PCRs in 25μl reactions containing 2 μl DNA template, 1U MyTaq polymerase (Bioline, Australia), 0.2 μM of each PCR primer, and 1x buffer. For amplification of *EF1alpha* and *16S* we performed PCR with 35 cycles at 52°C annealing, and for *Opsin* we used five cycles at 45°C annealing followed by 35 cycles at 50°C. PCR products were cleaned using 1U of Exonuclease I and Antarctic Phosphatase (New England Biolabs, Ipswich, Mass., USA) and incubating at 37 °C for 20 min followed by 10 min enzyme denaturation at 80 °C. Sequencing was conducted using the same forward and reverse primers used for PCR, by Macrogen Inc. (Seoul, Republic of Korea). We edited sequence data with CodonCode aligner, and aligned them using MAFFT in Geneious Prime 2019.2.1 (www.geneious.com, Kearse *et al*. 2012). We generated haplotype networks using the TCS algorithm (Clement *et al*. 2000) in PopART version 1.7 (http://popart.otago.ac.nz/) (Leigh and Bryant 2015).

### Mitogenome sequencing and assembly

The “Carbonaria group” of *Tetragonula* are now known to have several pseudogenes (or “numts”, nuclear insertions of mitochondrial genes) (Françoso *et al*. 2019; see also Franck *et al*. 20014, Brito *et al*. 2014). To ensure we were amplifying only the true *16S* gene and to provide a reference genome for each species for future work, we conducted whole genome sequencing on one individual of each of the three species and reconstructed the complete mitochondrial sequence. Sequencing libraries were made using the NebNext UltraII DNA kit for Illumina (New England Biolabs, Ipswich, Mass., USA), and PE150 sequencing conducted by Novogene (Beijing, China).

Sequences were checked with FastQC (Andrews, 2019), then we removed adaptors, and quality-trimmed the end of the sequences of all samples to q10 using bbduk from the bbtools package (version 36, Bushnell, 2019), with a kmer of 8. We binned the sequences by depth using a kmer-based approach in bbnorm (Bushnell, 2019). We put everything under 500x coverage into one file (this includes the nuclear genome, which had between 23x and 51x coverage for each species, Table 1), everything between 500x and 3,000x into another file (this is repetitive sequence and other artefacts below the depth of the mitochondria), everything over 3,000x coverage was placed in a third file, and this contained mitochondrial sequence and anything else with a depth over 3,000. This process excluded nuclear-mitochondrial pseudogenes and also reduced the amount of data going into subsequent steps. We then reconstructed mitogenomes by mapping reads to the mitogenome of the stingless bee *Melipona bicolor* (AF466146.2, Araujo & Arias 2019), followed by *de novo* assembly of the mapped reads, mapping and extension to the longest contigs, and then joining of contigs and manual verification of the final sequence by read-mapping (Geneious version 2019.2., Kearse *et al*. 2012). The final mitogenome assemblies were annotated using DOGMA (Wyman *et al*. 2004) and MITOS (Bernt *et al*. 2013), with reference to the *M. bicolor* mitogenome and manual editing in Geneious.

**Table 1.** Characteristics of the genomes and their draft assemblies for three species of *Tetragonula* stingless bees, including genome size, coverage, and % repeat, which were estimated with a kmer analysis in bbnorm. The mitogenome coverage was assessed by read mapping, and the assembly statistics (length, longest contig, n50, number of contigs) were calculated in Quast. The genome assemblies can be found at genbank under the provided bioproject and biosample IDs.

| **Species** | **Genome size estimate** | **Nuclear genome coverage** | **Mitogenome coverage** | **% repeat** | **Assembly length** | **Longest contig** | **N50** | **N contigs** | **BioProject** | **BioSample** |
| --- | --- | --- | --- | --- | --- | --- | --- | --- | --- | --- |
| *T. carbonaria* | 457,237,826 | 23x | 6,899 | 33% | 284,886,424 | 187,309 | 14,476 | 156,012 | PRJNA578948 | SAMN13088838 |
| *T. hockingsi* | 376,762,141 | 35x | 11,089 | 26% | 288,263,186 | 200,717 | 11,462 | 225,627 | PRJNA578994 | SAMN13089661 |
| *T. davenporti* | 342,107,567 | 38x | 26,907 | 17% | 288,842,417 | 271,029 | 18,924 | 72,464 | PRJNA578998 | SAMN13089823 |

We aligned these three new bee mitogenomes with three more unpublished *Tetragonula* mitogenomes and the *M. bicolor* mitogenome (AF466146.2) using MAFFT (Katoh and Standley 2013). The most appropriate evolutionary model for each gene was estimated using jModelTest 2.1.10 with the best model chosen based on Bayesian Information Criterion (BIC). We then constructed a bayesian tree with MrBayes (Huelsenbeck & Ronquist 2001).

### Nuclear genome assembly

To investigate further the low nuclear divergence suggested by our two nuclear genes (*Opsin* and *EF1alpha*), we considered the remaining genome-wide data generated from our Illumina sequencing. We took the lower genomic bin of the Illumina data (everything below 500x coverage based on kmer binning; Bushnell 2019), and assembled each species using SPADES (Bankevich *et al*. 2012), which was run with default settings, including the in-built error correction module. We assessed the assemblies and generated basic statistics using QUAST (Gurevich *et al*. 2013). We then took the 768bp *EF1alpha* PCR product and searched it against custom-blast databases for each genome. We did the same for *Opsin*. These larger sequences were aligned with MAFFT and a Bayesian tree was inferred from this alignment with MrBayes.

## Results

### Tests of hybridisation using microsatellites and snps

PCAs using both microsatellite and *snp* datasets separated samples into three non-overlapping clusters consistent with their species assignment (<5% missing data, Fig. 2A). The microsatellite PCA placed *T. davenporti* closer to *T. carbonaria* than to *T. hockingsi* (Fig. 2A), but the *snp* PCA indicated an equal distance between the three clusters (20% of the variance explained by each of the first two components, Fig. 2B). The loading plot of the *snp* data indicated that many of the 2,043 *snp* markers were contributing to the clustering in the PCA, rather than a few specific loci (Fig. 2C).

**Figure 2.**
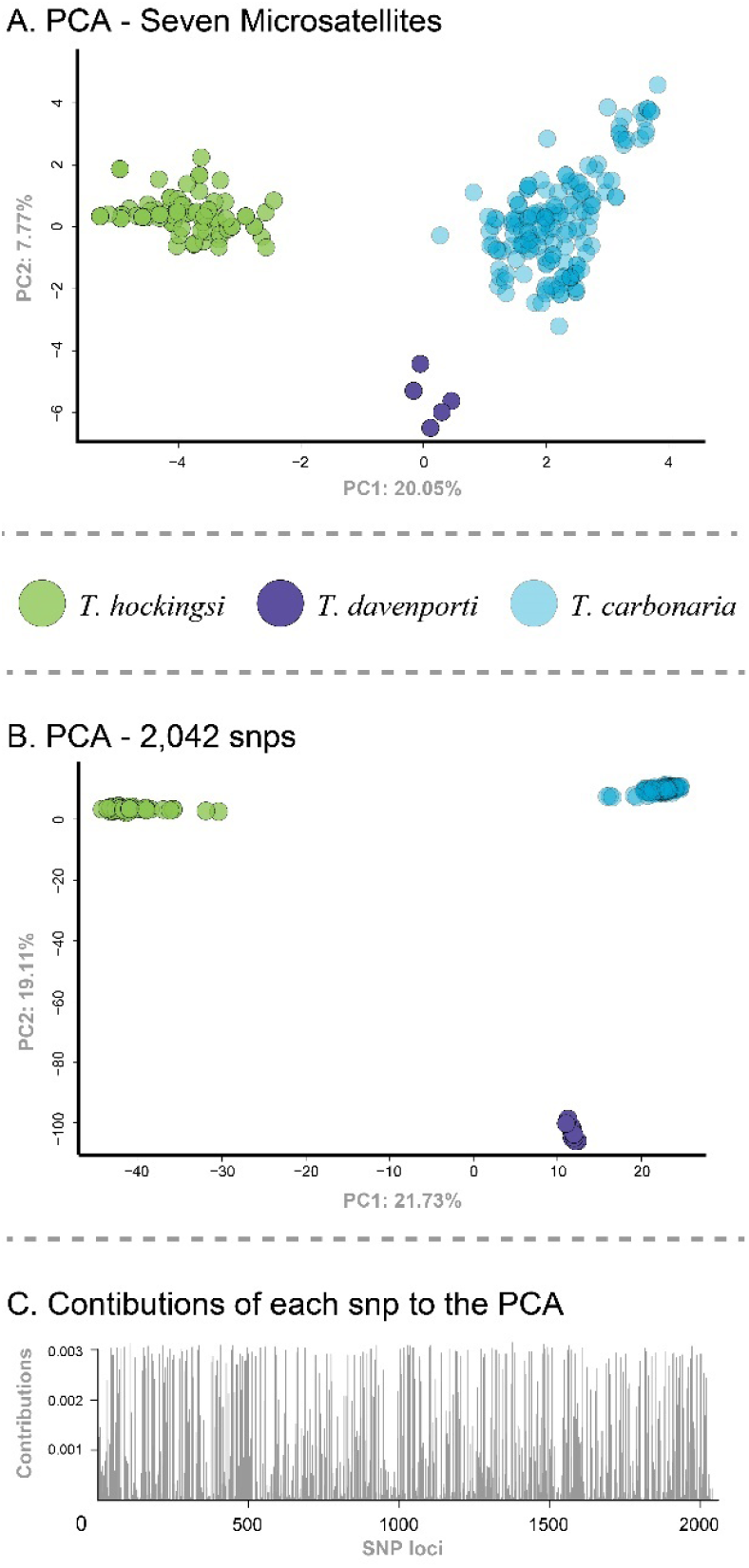
Principal Components Analysis (PCA) of genetic distance based on nuclear DNA markers. Plot showing principal components one and two based on the seven microsatellite markers (A), the same plot based on the 2,049 nuclear *snp* markers (B), and a loading plot showing the relative contributions of each of the 2,049 *snp* markers to the PCA (C).

*STRUCTURE* analyses using both microsatellite and *snp* datasets (N= 63 colonies, one worker per colony) returned no evidence of hybrids, with all individuals showing close to 100% assignment to a species’ cluster (Fig. 3). Notably, *STRUCTURE* results were sensitive to sibling-sampling and missing data. The same analyses using datasets comprising four workers per colony (akin to resampling the same individual multiple times), assigned four individuals (microsatellites) and two individuals (*snps*) at less than 90% posterior probability of belonging to their species’ cluster, thus identifying them as putative hybrids (Fig. 3). Likewise, subsampling to 20% missing data resulted in two *T. hockingsi* individuals being designated as hybrids with *T. davenporti*, but only with the microsatellite data. Missing data had no impact on the *snp* dataset, i.e. every individual was still assigned 100%.

**Figure 3.**
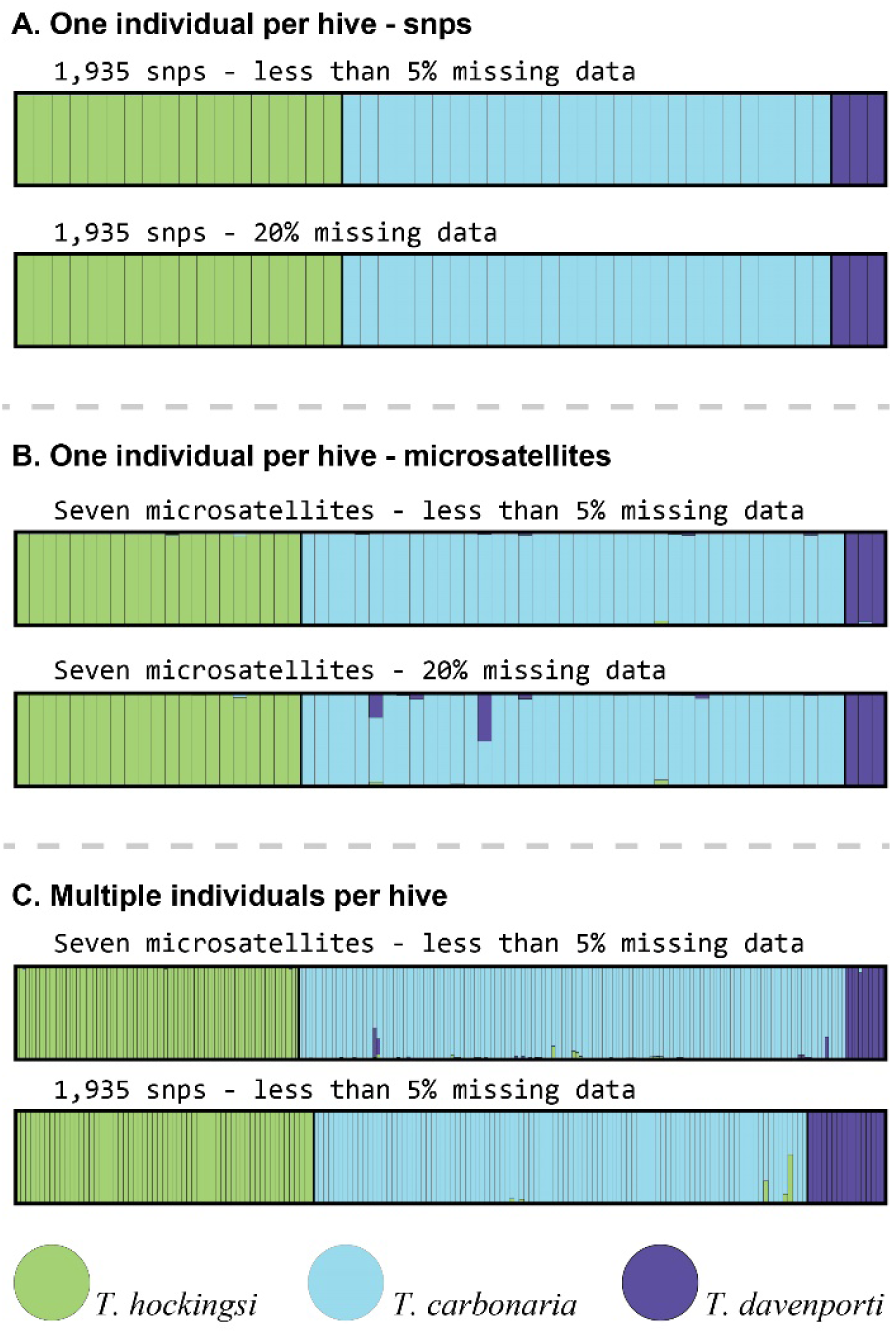
Structure plots assuming three hypothetical populations (K = 3), each bar represents an individual and the proportions of the three colours represent the posterior probability of that individual belonging to each of the three hypothetical (K) populations. Analyses were conducted for two *snp* datasets, one with 1,935 *snps* and less than 5% missing data, and one with the same number of *snps* with 20% missing data, using one individual per hive (A). The same analyses were conducted on the two datasets based on seven microsatellite loci and one individual per hive, one with less than 5% missing data and one with 20% missing data (B). The analyses were also run with multiple individuals per hive, mostly four but sometimes fewer, to examine the effect of including closely related siblings (C).

### Species-specificity of PCR-amplified gene regions

The mitochondrial gene *16S*, and the nuclear gene *Opsin*, each grouped samples into three haplotype clusters consistent with species assignment (Fig. 4), although pairwise divergence between species was greater for *16S*, with ∼6% between *T. carbonaria* and *T. hockingsi* based on 229 individuals representing all 63 colonies than for *Opsin*, with ∼0.5% difference between *T. carbonaria* and *T. hockingsi* (116 individuals representing all 63 colonies). The interspecific sequence difference in the *16S* fragment was considerably lower than that of the complete mitogenome (see below), because half of the amplified region was highly conserved (no variation). The nuclear gene *EF1alpha* had identical sequence for all samples (245 individuals representing all 63 colonies). No individuals showed mitochondrial-nuclear discordance between *Opsin* and *16S*. That is, if an individual had a *16S* sequence representative of *T. carbonaria* it also had an *Opsin* sequence representative of *T. carbonaria* (Fig. 4). However, one colony that was labelled *T. carbonaria* during field collections returned sequences and nuclear markers consistent with *T. hockingsi* and was assumed to be the result of either labelling error, or a recent interspecific colony takeover. Further, one colony suspected of being *T. davenporti* was genetically assigned to the *T. carbonaria* cluster, highlighting the difficulty of identifying them without genetics.

**Figure 4.**
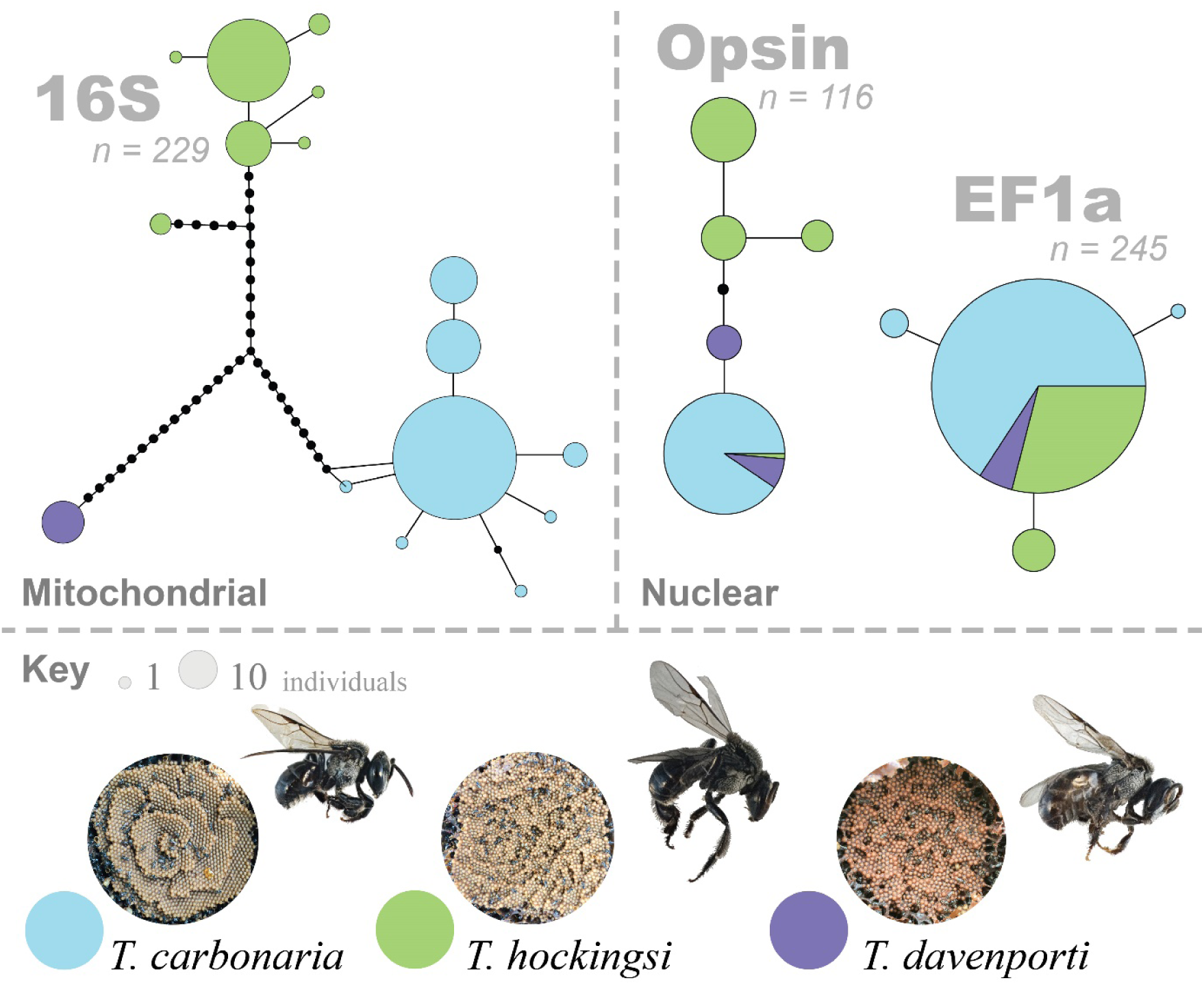
Haplotype networks for the mitochondrial *16S* gene, and nuclear *Opsin* and *EF1alpha* (top). Brood of the three species we investigate is illustrated as are representative bees of each of the three *Tetragonula* species in the key (bottom).

### Mitogenomes and genomes

We were unable to fully resolve the AT-rich control region of the mitochondria, but we did recover all mitochondrial genes. Our assemblies of the mitogenomes for these three species indicated 22.2% pairwise difference across *T. carbonaria* and *T. hockingsi*, 21.6% across *T. carbonaria* and *T. davenporti*, and 23.6% across *T. davenporti* and *T. hockingsi* (Fig. 5). Using these mitogenomes we were able to determine that the *16S* mitochondrial sequences were of mitochondrial origin and not nuclear pseudogenes. An obvious gene inversion was detected, relative to *M. bicolor* and all other bees for which mitogenome sequences are available (Supplementary Fig. 1) with *12S, 16S, NAD1, CYTB* and *NAD6* reversed and inserted in between *NAD3* and *NAD5*.

**Figure 5.**
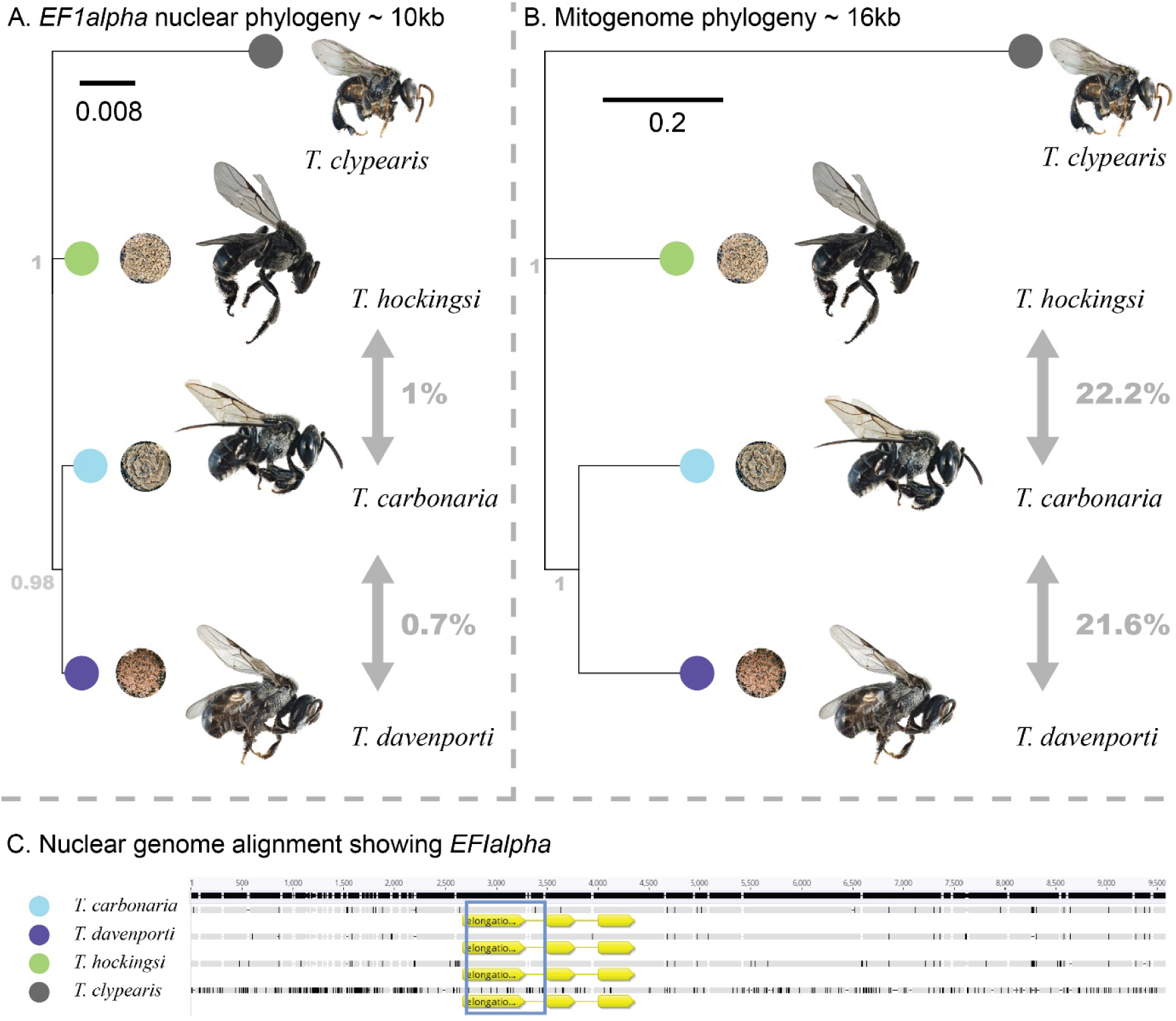
Bayesian tree of four Australia *Tetragonula* sp generated from a 10kb surrounding the nuclear *EF1alpha* gene (A). Bayesian tree constructed from whole mitogenomes (B). The nuclear genome alignment of *EF1alpha* and flanking regions; the blue rectangle represents the PCR product (C).

Nuclear genomes returned a mean *n50* of 14,954 across the three species (Table 1). From this sequencing, we investigated further the lack of interspecific difference at *EF1alpha*. We took the *EF1alpha* sequence and used it as a ‘bait’ to pull out the regions of the genome surrounding the 768bp PCR product. We managed to select at least 10kb in each species that aligned around the PCR product using MAFFT (Katoh and Standley 2013). Duplications were not present as only one hit was found in each assembly. Variation was detected around the *EF1alpha* locus and this separated the three species, based on the one individual of each that we sequenced. We thus confirmed that the pairwise difference across these three bee species was indeed extremely low in this region of the nuclear genome (0.7-1%, Fig. 5), relative to that detected in the mitochondrial genome. Similarly, we recovered 26kb around the *Opsin* gene, which returned the same low nucleotide difference between species (0.7-1%).

A Bayesian phylogenetic analysis based on either mitogenomes, or the flanking regions of *EF1alpha*, resolved the evolutionary relationships between the *Tetragonula* species in the same way, with *T. carbonaria* and *T. davenporti* diverging from each most recently, and *T. hockingsi* as their sister group (Fig 5).

## Discussion

### Testing for hybridisation

Hybridisation and gene flow are central to our understanding of species and speciation, but accurately detecting hybridisation depends on the markers and analyses used. In the case of Australia’s *Tetragonula* stingless bees, two previous studies concluded a low incidence of hybrid colonies in an area of sympatry between sister species (2-3% of all sampled colonies; Franck *et al*. 2004, Brito *et al*. 2014). In this study, we use multiple additional molecular markers to analyse a new sample of colonies from the same area of sympatry (South-East Queensland) and find all markers reliably segregate the three species, *T. carbonaria, T. hockingsi* and *T. davenporti*. We cannot discount rare occurrences of hybrid colonies based on 63 hive samples, but we can conclude that there is no gene flow between the species in this region beyond the F1 generation from our data.

The lack of gene flow between *Tetragonula* species indicates that these species either do not mate, or that interspecific mating produces inviable or infertile offspring. Beekeepers regularly keep multiple species in close proximity, providing opportunities for interspecific mating; for example, at one property that we sampled, colonies of all three species had resided together for at least nine years. In stingless bees, mating occurs on the wing close to the queen’s colony. Males typically aggregate out the front of a colony that is undergoing the re-queening process (i.e. has a virgin queen), presumably attracted by a pheromone (Vollet-Neto *et al*. 2018). Male aggregations may attract dozens to hundreds of unrelated males from the surrounding area (Vollet-Neto *et al*. 2018; Cameron *et al*. 2004). Virgin queens then fly into the aggregation and mate with just one male, before returning to the nest (Green & Oldroyd 2002). Even if aggregations sometimes contain males of more than one *Tetragonula* species, it may be that males are only sufficiently attracted to conspecific queens to attempt actual mating. Alternatively, queens may be crucial to the recognition process at the point a male makes contact with them, as in *Aphytis lingnanensis* parasitoids (Fernando and Walter 1997). Further, interspecific mating may occur, but produce inviable diploid brood due to genetic incompatibility. This latter scenario has been reported in sister species of honeybee that have come into sympatry recently (*Apis cerana* and *Apis mellifera*). Interspecific mating between these honeybees is common, but all hybrid offspring die at the early larval stage (Remnant *et al*. 2014; Gloag *et al*. 2017). In this case, mis-mated queens may be quickly replaced. For example, workers of at least one stingless bee species (*Scaptotrigona depilis*) will execute laying queens that have made inbred matings and are producing diploid male brood (Vollet-Neto *et al*. 2017). Clearly, further investigation into the mating of *Tetragonula* species is required to clarify the interactions between males and females in mixed species swarms.

Australian *Tetragonula* highlight two common reasons why it can be difficult to identify hybridisation between sister taxa. First, *Tetragonula* have nuclear pseudogenes of several common mitochondrial marker genes (Françoso *et al*. 2019; Franck *et al*. 2004). Such pseudogenes can complicate efforts to assess hybridisation through mito-nuclear discordance, because they may be mistaken for valid haplotypes that are shared across species and thus mistakenly taken as evidence of introgression. Even where researchers aim to avoid pseudogenes, by checking for stop codons or frame-shifting indels in their sequences, a subset of pseudogenes may still be missed. The reconstruction of whole mitogenomes from short read data, using a depth cut-off to avoid nuclear copies, as in this study, may be necessary to identify true mitochondrial genes. A further issue is that the pattern expected from mito-nuclear discordance (a sample returning one mitochondrial sequence and a different nuclear one) is the same as one gets when samples are mislabelled. Rates of mislabelling have been estimated at around 4% in published genetic datasets (Zych *et al*. 2017), so any cases of mito-nuclear discordance at these rates probably deserve additional scrutiny. We detected one instance of mislabelling across two of our datasets, by virtue of having conducted multiple genetic tests on the same individuals. Both pseudogenes and “mis-labelling introgression” have likely led to mistaken assignments of hybridisation in many taxa, and inflated perceptions of the extent to which species hybridise.

Second, where nuclear markers suggest low divergence between species, the interpretation of *STRUCTURE* analyses must take care to avoid biasing group assignment or over-interpreting structure plots (Lawson *et al*. 2018). In our dataset, the inclusion of four individuals per colony (full sisters) in a *STURCTURE* analysis led to the assignment of some individuals as putative hybrids. When this pseudo replication was removed, however, by including just one individual per colony, no hybrids were evident. Likewise, missing data had a large effect on the likelihood of inferring *Tetragonula* hybridisation in our dataset. Our original microsatellite dataset had low levels of missing data (<5%) and every individual was assigned at close to 100% into a species group. When we subsampled this dataset to 20% missing data, however, some samples appeared to be hybrids, indicating that *STRUCTURE* analyses can be sensitive to such levels of missing data. Notably, in this test on the effect of missing data, we subsampled our microsatellite data randomly across markers. In real datasets, microsatellites tend to go missing in a much less random way, generally with one or two loci most affected. The impact of missing microsatellite data is therefore likely to be even greater in practice than in our simulation. *STRUCTURE* assignments using microsatellites were the basis of hybrid identification in previous studies of Australian *Tetragonula* and may have been sensitive to these issues (Franck *et al*. 2004; Brito *et al*. 2014). In our study, *snp* data was much less sensitive to the effect of missing data and vastly outperformed microsatellite data to test for hybridisation. Indeed, in taxa where nuclear genome divergence is low, using many markers may be the only way to confidently assess hybridisation rates. In the absence of direct parentage analysis to assess hybridisation, such a multi-marker nuclear genome assay is the best approach to infer whether gene flow between species is ongoing (Twyford & Ennos 2011).

### Mitochondrial divergence between Tetragonula species

Our mitogenome data support previous studies indicating a high mitochondrial sequence divergence between species of the ‘Carbonaria complex’ (Cunningham *et al*. 2012; Francosco *et al*. 2019). This mitochondrial divergence contrasts with the low divergence we observed in two nuclear genes and their flanking regions, around 0.7-1% bp difference *vs*. 21-23% in the mitogenomes. We investigated the lack of variation in the *EF1alpha* PCR product further to see whether it was restricted to this 700bp fragment. We were able to recover the sequence flanking this region from our whole genome assemblies. These larger fragments did exhibit variation and recovered the same phylogeny as the mitogenomes. The discrepancy in divergence might indicate a particularly high mutation rate for mitochondria in this clade. The ratio of mitochondrial to nuclear mutation rate varies substantially across metazoans (from at least 0.8x to 40x, Allio *et al*. 2017). A mutation rate in *Tetragonula* mitochondria that was around 20 times that of their nuclear genome would be within the range reported for insects, albeit at the high end (Allio *et al*. 2017; Francosco *et al*. 2019). Several other Hymenoptera have high rates of mitochondrial divergence, compared to other insects (Castro *et al*. 2002; Kaltenpoth 2012). This pattern of mito-nuclear substitution rate difference could also be interpreted as being evidence for a relatively deep divergence time for these species (evidenced in the divergent mitochondria) that was later followed by a period of nuclear gene flow, erasing nuclear divergence. Under this scenario, past gene flow between diverged populations must have occurred in a similar fashion between all three species, because they share a common ratio of mitochondrial to nuclear divergence. Distinguishing between these two alternatives requires a better understanding of rate variation in mitochondrial and nuclear genomes of stingless bees relative to other Hymenoptera, and population genomic studies to elucidate the evolutionary history of this group.

*Tetragonula davenporti* remains the most mysterious of the three *Tetragonula* species considered in this study. It has been confirmed from only one property in the Gold Coast hinterland (originally sourced from forested areas nearby) and one wild specimen from the Queensland-New South Wales border (R. Gloag, unpublished data). It is essentially cryptic in morphology with *T. carbonaria* and in nest structure with *T. hockingsi*. Nevertheless, the distinctive mitogenome of *T. davenporti* reported here lends strong support to its status as a valid species. The natural range of *T. davenporti* is currently unknown, and the difficulty in identifying it morphologically to species means that its presence among managed colonies is also unknown. South-east Queensland is an area of high endemicity, so it may well be endemic in this area.

This study investigated managed colonies mostly from Brisbane and SE Queensland because this is where hybridisation is most likely between these three species and has been reported previously. However, *T. carbonaria* and *T. hockingsi* also co-occur in parts of Far North Queensland, and *T. hockingsi* is increasingly being transported by beekeepers into more southern parts of *T. carbonari*a’s range (pers. obs. TJS). Examining wild populations of this genus may reveal additional species within the ‘*carbonaria* complex’. Comparing wild populations to those that have been transported anthropogenically will provide further insight into the impact of human movement on the evolution of this group. Finally, we hope the genomic resources developed here will enable further investigation of this fascinating group of social bees.

## Acknowledgements

We would like to acknowledge all the beekeepers and hive owners who donated bees to this study, especially Tim Heard, Peter Davenport and Alan Lashmar. We thank the University of Queensland for financial support.

## Data Availability

The PCR amplicon data can be found on Genbank; acessions MN659448 - MN659676 for 16S, MN659677 - MN659921 for EF1alpha and MN659922 - MN660037 for Opsin.

Individuals used for whole genome sequencing have NCBI Biosample numbers as shown in Table 1. Illumina data has been deposited at the Sequence Read Archive (SRA), *T. carbonaria* = SRX7089578, *T. hockingsi* = SRX7104558, *T. davenporti* = SRX7104558. Genome assemblies can be found on Genbank, *T. carbonaria* = GCA_010645115.1, *T. hockingsi* = GCA_010645185.1, *T. davenporti* = GCA_010645165.1. These can all be found on the University of Queensland espace as well (Hereward et al. 2020, doi: https://doi.org/10.14264/84de1f8).

Microsatellite data, raw *snp* data, vcf files, R scripts, additional genome assembly data, and locations and dates of samples can be found on the University of Queensland espace (Hereward et al. 2020, doi: https://doi.org/10.14264/84de1f8) The locations are only accurate to around 1km to protect the privacy of beekeepers.

## Author Contributions

All authors conceived the project, TJS obtained samples, JPH conducted molecular lab work, JPH and DRB analysed the genetic data. All authors wrote the manuscript.

## Supplementary data

Supplementary Figure 1. Mauve alignment of the mitogenomes showing the inversion between *Tetragonula* and other bees. Supplementary Figure 2. PCA plot of the *snp* data highlighting the samples from Rockhampton (Queensland), Newcastle, and Sydney (both New South Wales). Supplementary Data 1. Protocol for the genotyping by sequencing assay. Supplementary Table 1. Spreadsheet of the GBS adaptors.

