## Supplementary material for "Tests of hybridisation in *Tetragonula* stingless bees using multiple genetic markers": Supplemetary Data 1

**Description**

This is the GBS method we have optimised based on [Elshire 2011](https://journals.plos.org/plosone/article?id=10.1371/journal.pone.0019379), [Poland 2012](https://journals.plos.org/plosone/article?id=10.1371/journal.pone.0032253) and [Petersen 2012](https://journals.plos.org/plosone/article?id=10.1371/journal.pone.0037135), with golay barcodes from [Caporaso 2012](https://www.nature.com/articles/ismej20128). We usually run 300 samples per lane and get good results from a variety of different plant species. Currently all the Fwd barcodes are the same length so we still need to spike in 20% genomic DNA to keep the sequencer happy when it gets to the cut-site, future modifications will add a second plate of forward adaptors with additional barcode bases to remove this requirement.

**What you need**

1. Adaptors and primers – see google sheet: <https://docs.google.com/spreadsheets/d/1DRM2VCz8muT_Xv3Fk9vWeCpoLeHauj064wu1QXOEE7A/edit?usp=sharing>
2. PstI - [available from NEB](https://www.nebiolabs.com.au/products/r3140-psti-hf#Product%20Information)
3. MspI - [available from NEB](https://www.nebiolabs.com.au/products/r0106-mspi#Product%20Information)
4. Cutsmart Buffer - [available from NEB](https://www.nebiolabs.com.au/products/b7204-cutsmart-buffer#Product%20Information)
5. T4 DNA Ligase - [available from NEB](https://www.nebiolabs.com.au/products/m0202-t4-dna-ligase#Product%20Information)
6. 10mM ATP - [available from NEB](https://www.nebiolabs.com.au/products/p0756-adenosine-5-triphosphate-atp#Product%20Information)
7. Phusion Taq - [available from NEB](https://www.nebiolabs.com.au/products/e0553-phusion-high-fidelity-pcr-kit#Product%20Information)
8. Clean up beads - we use [PCRcleanDX](https://alinebiosciences.com/)

**Protocol**

**Anneal Adaptors**

Recipe for 5x Annealing Buffer (50mM Tris, 250mM NaCl, 5mM EDTA) to make 50ml:

| 1M TRIS-Hcl ph8 | 2.5ml |
| --- | --- |
| 0.5M EDTA | 500µl |
| 1M NaCl | 25ml |
| make up to 50ml with clean H_2_O |  |

Refer to adaptor sheet for the adaptors to order, we have 96 forward adaptors based on [Poland 2012](https://journals.plos.org/plosone/article?id=10.1371/journal.pone.0032253) with 12bp Golay barcodes from [Caporaso 2012](https://www.nature.com/articles/ismej20128), and then use 12 reverse adaptors based on [Petersen 2012](https://journals.plos.org/plosone/article?id=10.1371/journal.pone.0037135) that allow indexing, with variable length MTC codes to improve the quality of the reverse sequencing read.

1. To make 40uM of each adaptor add 40ul Fwd adapter oligo, 40ul Rev adapter oligo and 20ul of 5x annealing buffer.
2. PCR program GBS barcode anneal - incubate at 97.5^o^C for 2.5 minutes, and then cool at a rate of not greater than 3^o^C per minute until the solution reaches a temperature of 21^o^C. Hold at 4^o^C
3. Dilute down the adaptors - Fwd adaptors diluted to 5ng/µl (10x) then 0.5ng/µl (working dilution). Rev adaptors kept at 40µM

**Normalise DNA**

Prepare 100-1000ng DNA in 40µl. Normalise using pico green. If there is variation in the integrity (degradation) pool by integrity when pooling for the pippin prep so that the normalisation of pools evens it out. If you don't do this, you may get 2x reads for the high integrity samples. We don't normalise again during the protocol, so it is important to get it right at the start.

**Double Digest**

**Recipe:**

| **Ingredient** | **1 sample** | **96 well plate** |
| --- | --- | --- |
| PstI | 0.5µl | 55µl |
| MspI | 0.5µl | 55µl |
| Cutsmart Buffer | 5µl | 550µl |
| Water | 4µl | 440µl |

1. Dispense 10µl per well using the step pipettor.
2. PCR free PCR machine DoubleDigest 3hrs @ 37^O^C, 20min @ 65^O^C, 4^O^C forever.
3. Go directly to ligation

**Adaptor Ligation**

**Ligation mix recipe:**

| **Ingredient** | **1 sample** | **96 well plate** |
| --- | --- | --- |
| 10mM ATP | 6.5µl | 715µl |
| T4 DNA ligase | 0.5µl | 55µl |
| Cutsmart Buffer | 1.5µl | 165µl |
| Water | 1.5µl | 165µl |

1. Add 4µl of P1 barcode adaptor (0.5ng/µl) and 1µl of P2 adaptor to each sample with multichannel pipette and filtered tips taking great care not to mix up the adaptors - this is a critical point where any mistakes will result in "accidental introgression"
2. Dispense 10µl per well using the step pipettor.
3. PCR free PCR machine GBSLigate 3 hrs @ 22^O^C, 20min @ 65^O^C, 4^O^C forever.

**Pooling and size selection**

1. Using multichannel pool all 96 samples into one disposable reservoir, mix using P1000 pipette.
2. Keep 2 x 1.5ml as a safety stop (freeze for later just in case).
3. Add 500µl of pooled GBS, 500µl of 4M GuHCL and 500ul of 100% EtoH to each of six eppies.
4. Mix, add contents of all six eppies into one spin column per plate (750µl at a time). Spin down for 1 min and repeat until all GBS has been bound to the silica columns.
5. Proceed to one wash with 750µl wash buffer, add buffer and then spin for 1 min, discard eluate and then spin for 2 mins to dry column.
6. Add 100µl of 10mM Tris to each column - incubate 1 min then spin into collection eppie.
7. Take 40µl of this and size select 300-400bp on the pippin prep, the other 60µl is kept as a second safety stop (freeze like the other).

Note: Pippin prep manual says use 30ul, and add 10ul of the dye/buffer mix, but you can fit 40+10=50ul total in there.

**PCR amplification**

**PCR Recipe:**

| **Ingredient** | **1 sample** | **11 samples** |
| --- | --- | --- |
| Phusion 5x HF | 6µl | 66µl |
| 10mM DNTP | 0.6µl | 66µl |
| Illumina Fwd PE | 1.2µl | 13.2µl |
| Rev Index primer | 1.2µl | 13.2µl |
| Taq | 0.24µl | 2.64µl |
| Water | 0.76µl | 8.36µl |

1. Take 2x 20µl aliquots removed from pippin prep, add 10µl mmix as above for 2 x 30µl reactions per index
2. PCR machine DDRadAmplify. 98^O^C for 2m, then 15 cycles of 98^O^C for 30s, 62^O^C for 20s, 72 ^O^C for 30s, final extension of 72^O^C for 5m.

Note: We usually pool three 96well plates together, and use one index for each, so Index 2, Index 6, and Index 12 as recommended to balance the index reads. Do not try to reduce the number of cycles until you are sure everything is working well.

**Post PCR Pooling**

1. Check each pool on Shimadzu (3µl library and 6µl water).
2. Check on Qubit (1µl).
3. From shimadzu calculate the proportion that is library = library peak/(library peak + other (adaptor) peaks)
4. Multiply Qubit reading by shimadzu proportion.
5. Make equimolar pool of indexes using the above figure and including 20% spike-in of genomic shotgun or other random illumina library.

**Bead Clean**

We do one 0.8X bead clean at the end to remove adaptors.

1. Allow beads to come to room temperature and then vortex them very well to resuspend.
2. For every 100µl of pooled library add 0.8X = 80µl of beads (ampure or equivalent, we use PCRcleanDX). Mix well by pipetting.
3. Incubate on bench for 10m.
4. Place the tube on an appropriate magnetic stand to separate the beads from the supernatant. If necessary, quickly spin the sample to collect the liquid from the sides of the tube before placing on the magnetic stand.
5. After 5 minutes (or when the solution is clear), carefully remove and discard the supernatant. Be careful not to disturb the beads that contain DNA targets (do not discard beads).
6. Add 200µl of freshly prepared 80% ethanol to the tube while in the magnetic stand. Incubate at room temperature for 30 seconds, and then carefully remove and discard the supernatant. Be careful not to disturb the beads that contain DNA targets.
7. Repeat the ethanol wash for two washes total. If necessary, briefly spin the tube, place back on the magnet and remove traces of ethanol with a p10 pipette tip.
8. Air dry the beads for up to 5 minutes while the tube is on the magnetic stand with the lid open. It is important not to over dry the beads - it can result in lower recovery of DNA target. Elute the samples when the beads are still dark brown and glossy looking, but when all visible liquid has evaporated. When the beads turn lighter brown and start to crack, they are too dry.
9. Remove the tube from the magnetic stand. Elute the DNA target from the beads by adding 22µl of 10mM Tris.
10. Mix well by pipetting up and down 10 times, or on a vortex mixer. Incubate for at least 2 minutes at room temperature. If necessary, quickly spin the sample to collect the liquid from the sides of the tube or plate wells before placing back on the magnetic stand.
11. Place the tube on the magnetic stand. After 5 minutes (or when the solution is clear), transfer 20µl to a new PCR tube, (optional) take 1µl of the remaining liquid and check on the Qubit, the pooled library is now ready to ship for sequencing. Either lyophilise and ship dry, or store at -20^o^C and ship on blue ice (ice bricks) that have spent the weekend in the -80^o^C freezer.

**James Hereward**

School of Biological Sciences
The University of Queensland


https://researchers.uq.edu.au/researcher/1855
