## Supplementary figures and images for "Tests of hybridisation in *Tetragonula* stingless bees using multiple genetic markers"

### Supplementary Figure 1

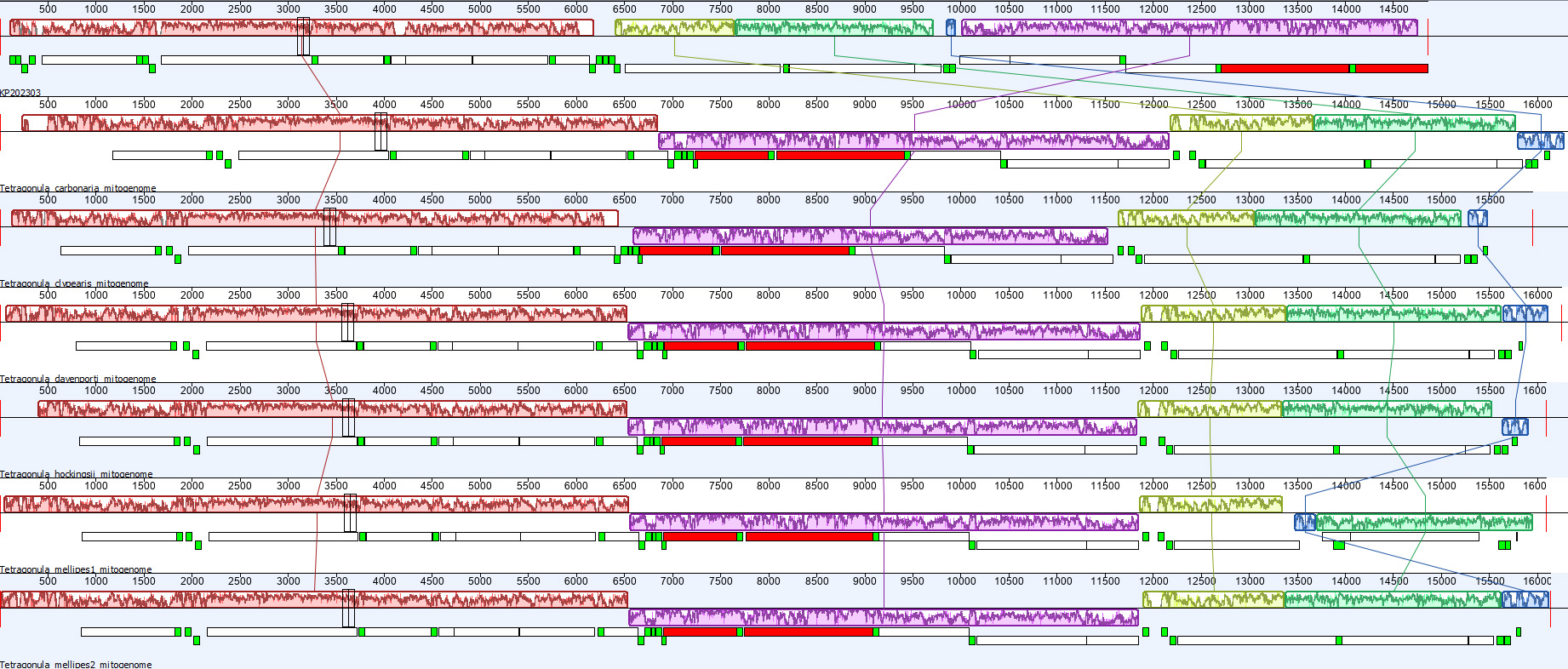

### Supplementary Figure 2

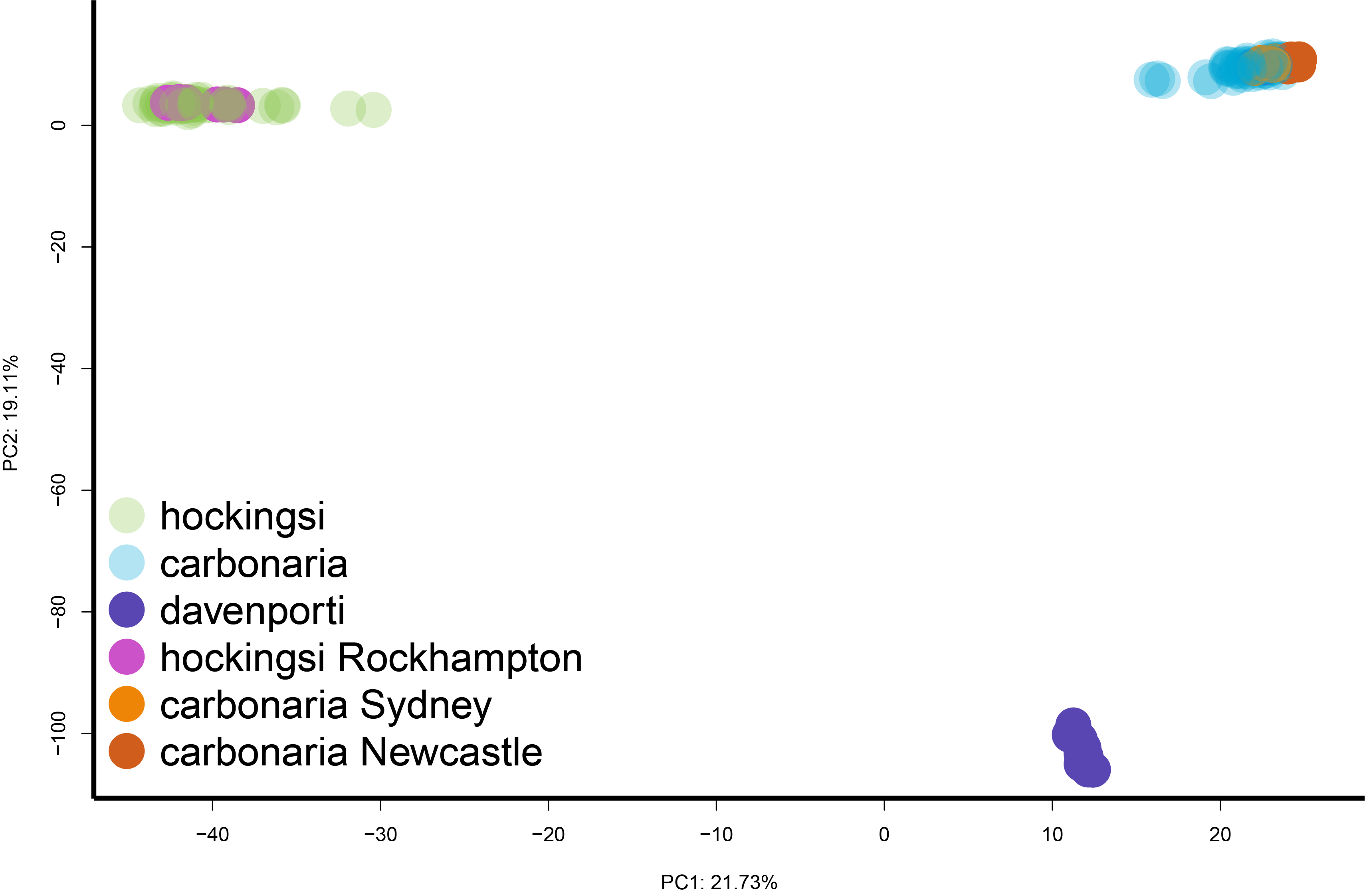
